# Chronic loss of STAG2 leads to altered chromatin structure contributing to de-regulated transcription in AML

**DOI:** 10.1101/856195

**Authors:** James S Smith, Stephanie G Craig, Fabio G Liberante, Katrina M Lappin, Clare M Crean, Simon S McDade, Alexander Thompson, Ken I Mills, Kienan I Savage

## Abstract

The cohesin complex plays a major role in folding the human genome into 3D structural domains. Mutations in members of the cohesin complex are known early drivers of myelodysplastic syndromes (MDS) and acute myeloid leukaemia (AML), with *STAG2* the most frequently mutated complex member. Here we use functional genomics to investigate the impact of chronic STAG2 loss on three-dimensional genome structure and transcriptional programming in a clinically relevant model of chronic STAG2 loss. The chronic loss of STAG2 led to loss of smaller loop domains and the maintenance/formation of large domains which in turn led to altered genome compartmentalisation. These changes in genome structure were linked with altered gene expression, including deregulation of the HOXA locus and the MAPK signalling pathway, which may contribute to disease development and response to therapy.

**AUTHOR SUMMARY:** Acute myeloid leukaemia (AML) and myelodysplastic syndromes (MDS) are clonal malignant diseases that affect the myeloid blood cell lineage. Around 40 different mutations have been identified as being associated with MDS and AML; several mutations effect genes in the cohesin complex particularly STAG2. STAG2 is an X-linked gene that is pivotal to the cohesin complex and the mutations result in a truncated gene and loss of function. We have introduced a clinically relevant truncating mutation into an isogeneic AML model. This has shown that changes in the sizes of loop and domains formed in the genome resulting in appropriate gene compartmentalisations. The associated changes in gene transcription has resulted in deregulation of HOX genes, essential for development and differentiation, and in the MAPK signalling pathway that could a therapeutic target.

## INTRODUCTION

Acute Myeloid Leukaemia (AML) is a highly clonal disease characterised by the rapid expansion of differentiation-blocked myeloid precursor cells, resulting in defective haematopoiesis, and eventually, bone marrow failure [1]. In recent years, large-scale sequencing studies have identified a plethora of mutations within patient derived AML cells, expanding our understanding of the genomic landscape, and highlighting the complexity of the varying subtypes of this disease [2]. One of the emerging genomic disease subgroups involves mutations within chromatin/spliceosome related genes; including members of the cohesin complex. Approximately 11% of patients diagnosed with a myeloid malignancy, including AML, Myelodysplastic syndrome (MDS) or Myeloproliferative neoplasm (MPN), have been shown to harbour a mutation within a member of the cohesin complex, with many more showing significantly reduced expression of the complex members [3–5]. Mutations within cohesin complex genes have also been identified in clinically normal and aging individuals following sequencing analysis of large populations, indicating that these mutations are early events in leukaemogenesis, leading to the identification of clonal haematopoiesis of indeterminate potential (CHIP) [6].

The cohesin complex is an evolutionarily conserved multimeric protein complex consisting of SMC1A, SMC3, RAD21 and either STAG1 or STAG2. The complex plays a pivotal role in mitosis through sister chromatid cohesion, as well as in homologous recombination mediated DNA double strand break repair [7] and has been suggested to be involved in long-range interactions between *cis* regulatory elements of the genome [8].

The most commonly mutated gene within the cohesin complex is STAG2, with ∼90% of STAG2 mutations resulting in the introduction of premature stop codons likely to lead to a loss of function [5]. The impact of loss of function STAG2 mutations on cohesin function has yet to be fully elucidated. Cohesin and the CCCTC binding factor (CTCF) have been described as master weavers of the genome [9], with a key role in regulating the 3D architecture of the human genome. CTCF and cohesin have been shown to co-localise heavily throughout the genome, separating regions of active and repressive chromatin marks regulating gene expression [9, 10].

In this study, the impact of a clinically relevant STAG2 mutation on 3D genome architecture, global and local gene expression and therapeutic potential was investigated.

## RESULTS

### GENERATION OF AN ISOGENIC ΔSTAG2 CELL MODEL

Given that the most common mutations within STAG2 result in the introduction of a premature stop codon early in the STAG2 coding region (including a mixture of frame shifting indels and stop gains)[3], we utilized a CRISPR/Cas9 system to target the *STAG2* coding region within the OCI-AML3 cell line. We selected the OCI-AML3 line, as it was derived from normal karyotype (NK) patients with NPM1c+ (Type A) and DNTM3A mutations, which have been shown to co-occur with STAG2 mutations in patient samples. We targeted the commonly mutated exon 20 of *STAG2* (ensuring all possible isoforms were affected), introducing a hemizygous 11 bp indel (STAG2 is encoded within the X-chromosome and is thus hemizygous in this male cell line) in our edited cell line (ΔSTAG2), resulting in the introduction of a premature stop codon 11 bp downstream of the sgRNA target site **(Figure S1A)**. Analysis of STAG2 mRNA highlighted a ∼7 fold reduction in expression compared to the *STAG2*-WT counterpart indicating that the mutant mRNA is degraded through non-sense mediated decay **(Figure S1B)**. When examining the mRNA expression of *STAG1* it was interesting to observe an increase of ∼1.5 fold relative to the *STAG2*-WT counterpart (**Figure S1C**). The other members of the cohesin complex (*RAD21*, *SMC1A* & *SMC3*) exhibited very similar patterns of expression at the mRNA level (**Figure S1C**). In keeping with this, STAG2 protein expression was completely abolished in this model **(Figure S1D)**. Additionally, when examining the expression of other members of the cohesin complex, including STAG1, similar protein levels were observed in both the ΔSTAG2 and STAG2-WT cells **(Figure S1D)**.

### STAG1 COMPENSATES FOR LOSS OF STAG2 CHROMATIN BINDING

Chromatin immunoprecipitation for STAG1, STAG2 and CTCF followed by deep sequencing (ChIP-Seq) was carried out to confirm the loss of STAG2 chromatin binding and to assess the chromatin binding of other core members of the cohesin complex in the absence of STAG2 (**Figure 1A**).

**Figure 1:**
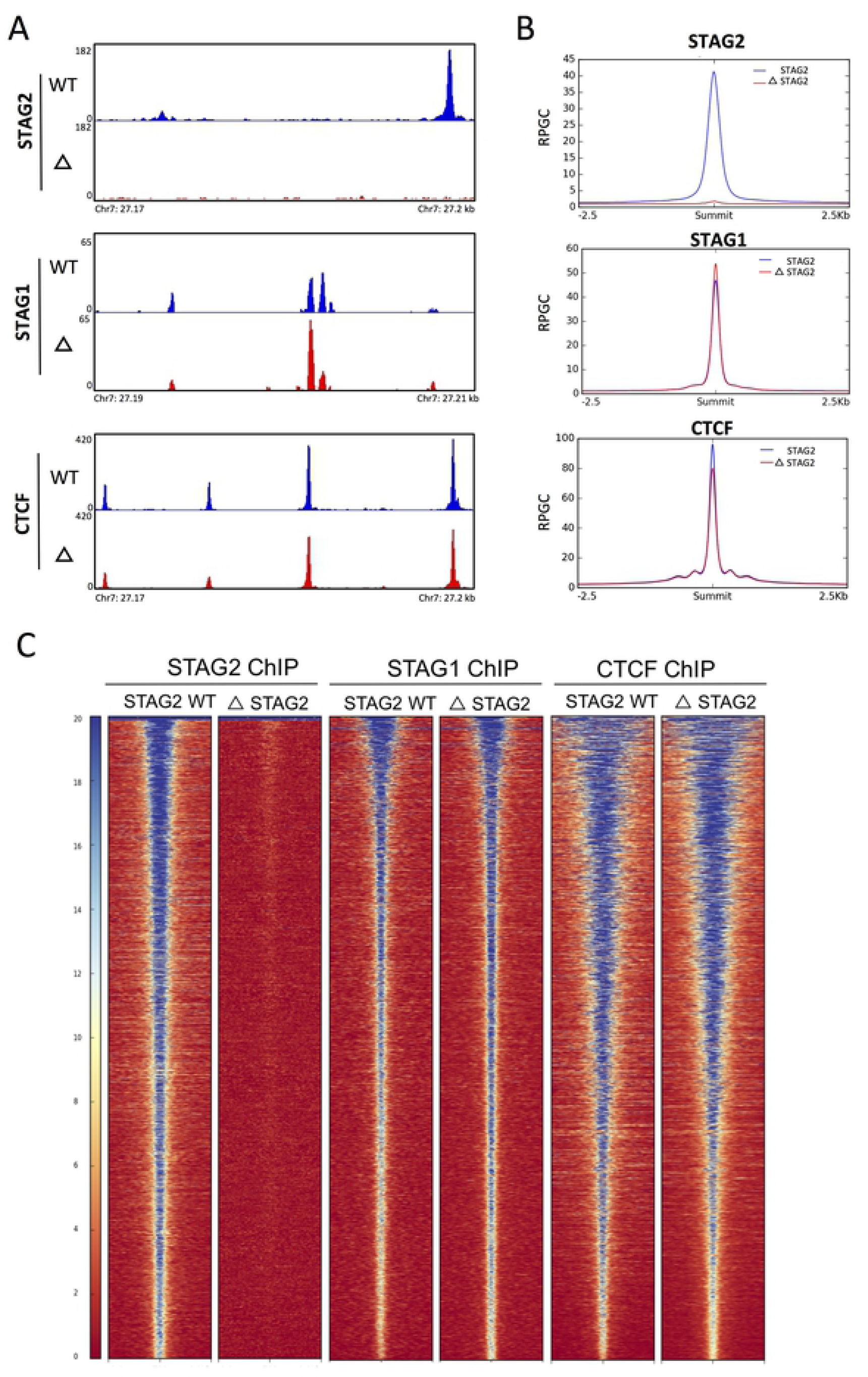
Cohesin binding patterns in the absence of STAG2 highlight STAG1 compensation. A) Normalised BigWig binding profiles generated from STAG2, STAG1 and CTCF ChIP-Seq, in STAG2-WT and ΔSTAG2 cells viewed in IGV. Each region shown, represents the typical binding pattern observed in these cells, for each protein throughout the genome and show the complete loss of functional STAG2, slightly increased binding STAG1 and slightly decreased binding of CTCF in the ΔSTAG2 cells, B) Consensus binding peak lists were generated for STAG2, STAG1 and CTCF from ChIP-Seq carried out in STAG2-WT cells. These lists were then used to plot the genome-wide average binding profiles of STAG2, STAG1 and CTCF at these consensus sites in STAG2-WT and ΔSTAG2 cells. Average binding profiles are normalised to total reads per genome content, centred on peak summits. C) Heatmap profiles of binding patterns for STAG2, STAG1 and CTCF in both STAG2-WT and ΔSTAG2 cells across the genome.

As expected, STAG2 binding was completely abrogated in the ΔSTAG2 cells, whilst STAG1 appeared to have slightly stronger binding profile in the ΔSTAG2 cells compared to STAG2-WT cells (**Figure 1B**). Conversely, the CTCF binding profile was slightly weaker in the ΔSTAG2 cells (**Figure 1B**).

To assess this at an individual binding region level, genome-wide signal density plots for STAG2, STAG1 and CTCF were generated using peak summit locations called in the STAG2-WT cells for each protein (**Figure 1C**). A total of 34,670 STAG2 binding peaks were identified in STAG2-WT cells, with none identified in the ΔSTAG2 cells, again confirming the complete loss of functional STAG2 in these cells. Analysis of STAG1 binding revealed a ∼35% increase in the number of STAG1 binding peaks (from 15,034 to 22,955) in the STAG2-WT and ΔSTAG2 cells respectively. Many of these additional STAG1 binding sites corresponded to canonical STAG2 binding sites suggesting a degree of compensation by STAG1 in the absence of STAG2. To confirm this, STAG1 peak calls from the STAG2-WT cells were intersected with STAG1 binding sites from the ΔSTAG2 cells. This showed that ∼96% (14,375/15,034) of the canonical STAG1 binding peaks were maintained in the ΔSTAG2 cells but also identified an additional 8, 580 new STAG1 binding sites, only present in the ΔSTAG2 cells **(Figure S2)**. To test if these new STAG1 binding peaks represented regions previously occupied by STAG2, we intersected the locations of these binding peaks with the STAG2 binding peaks called in STAG2-WT cells. Of the 8,580 new STAG1 sites, ∼86% (7,392) overlapped with identified sites in STAG2-WT cells; these also coincided with 8046 (94%) and 7947 (92%) CTCF sites in the STAG2-WT and ΔSTAG2 cells respectively.

This suggests that the increased STAG1 binding observed in the ΔSTAG2 cells has partially, but not completely, compensated for loss of STAG2 and was predominantly associated with CTCF binding sites.

### CHRONIC LOSS OF STAG2 LEADS TO ALTERED 3D GENOME ARCHITECTURE

3D chromatin structure is organised into loops and domains, which are anchored at convergent CTCF binding sites brought together through loop extrusion facilitated by cohesin[11]. Therefore, to examine 3D architecture, HiChIP analysis [12] targeting the CTCF to identify the CTCF/cohesin bound chromatin interaction sites present in both cell populations was undertaken.

Topologically associated domains (TAD’s) were identified using the Juicer Tools Arrowhead algorithm at both 10 kb and 25 kb resolutions, retaining unique domain calls from both resolutions to generate a consensus domain list [13]. This identified 5,014 and 622 domains in STAG2-WT and ΔSTAG2 cells respectively. However, despite the reduction in domains called in the ΔSTAG2 cells, the characteristic square appearance of a significant number of TAD’s was apparent when visually assessing contact maps generated from the ΔSTAG2 cells, but with significantly reduced signal **(Figure 2C)**. This indicates that some signal was present and visually observable, but was below the call detection threshold of the Juicer Arrowhead detection software. Nonetheless, far fewer TADs were detected in the ΔSTAG2 cells, suggesting that this may occur due to fewer contacts between CTCF anchor points.

**Figure 2:**
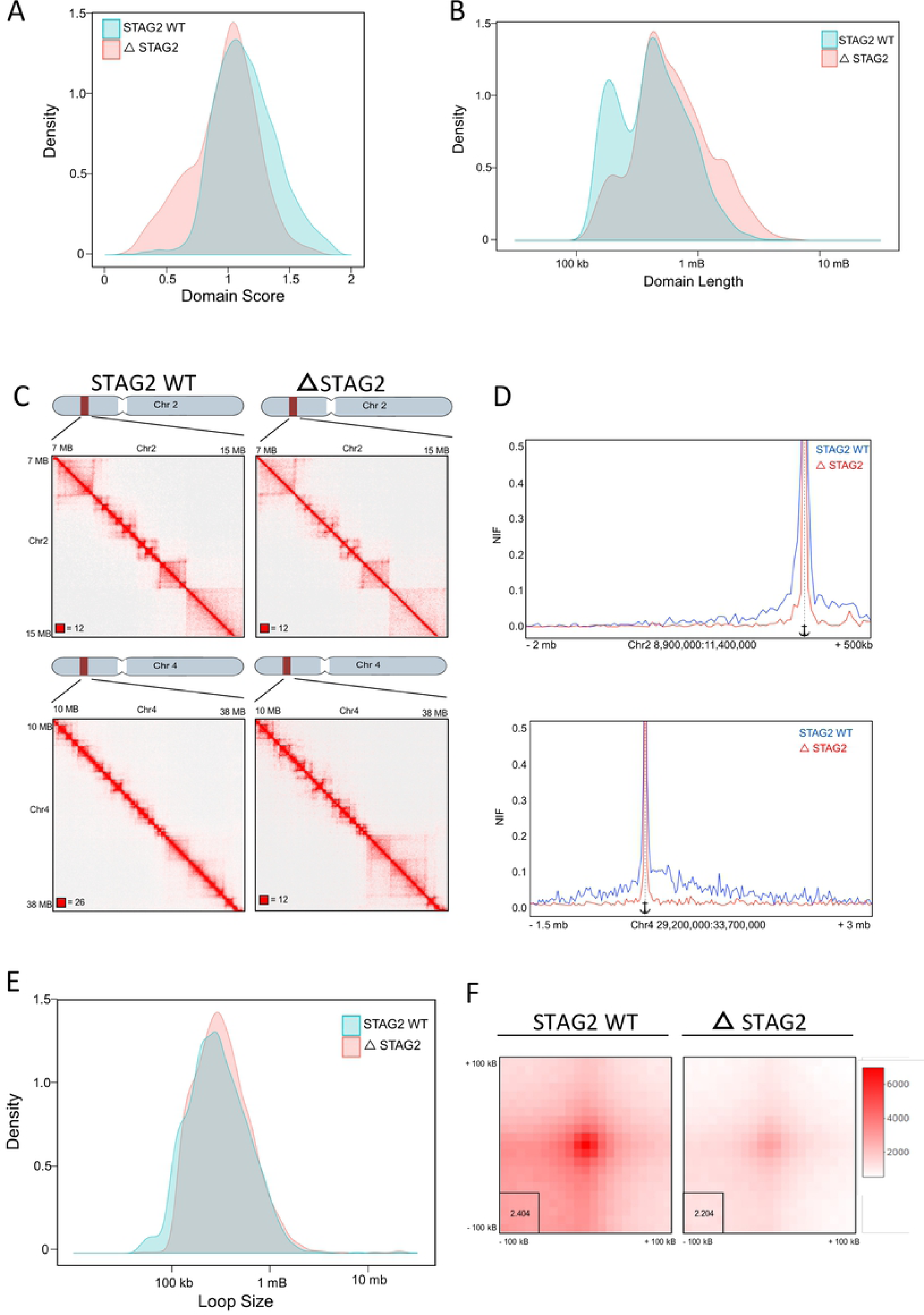
Loss of STAG2 alters domain architecture and interaction frequency of the 3D genome. A) Density plot showing the distribution of domain scores generated using the Arrowhead algorithm on CTCF HiChIP data generated from STAG2-WT and Δ STAG2 cells indicating reduced domain interaction frequency in ΔSTAG2 cells compared to STAG2-WT cells. B) Density plot showing distribution of domain lengths called in the STAG2-WT and ΔSTAG2 cells demonstrating loss of smaller domain interactions, maintenance of intermediate size interactions and increased frequency of larger domain interactions in ΔSTAG2 cells compared to STAG2-WT cells. C) Representative HiChIP Contact matrixes displaying interactions covering an 8 Mb region of chromosome 2 and a 28 Mb region of chromosome 4, highlighting the reduced interaction frequency of smaller loop domains, maintenance of intermediate sized domains and formation of larger domains in ΔSTAG2 cells compared to STAG2-WT cells. D) Virtual 4C (V4c) plot displaying interactions anchored at a domain boundary region 11 Mb into chromosome 2 and 30.7 Mb into chromosome 4 demonstrating significantly reduced interaction frequency with this anchor point in ΔSTAG2 cells compared to STAG2-WT cells. E) Density plot showing distribution of loop lengths called in the STAG2-WT and Δ STAG2 cells, indicating the formation of similar sized loops in both STAG2-WT and ΔSTAG2 cells. F) Aggregate Peak Analysis plots using loops called in the STAG2-WT cells as inputs, scaled internally to indicating significantly reduced loop signal intensity in Δ STAG2 cells in comparison to STAG2-WT cells.

The Juicer Arrowhead software[13] generates scores that serve as a measure of likelihood that the boundaries of the domains called are actual domain corners, with interactions likely to occur within the domain. We found that domains called in the STAG2-WT cells scored on average 1.14 (range 0.11 – 1.88) whilst in the ΔSTAG2 cells the score was reduced to an average of 0.96 (range 0.19 – 1.69) **(Figure 2A)**. The decrease in average score reflected the decreased frequency of domain forming interactions observed in the ΔSTAG2 cells.

Additionally, TADs in the STAG2-WT cells, ranged in size from 120 kb to 5.8 Mb (median 425 kb) with 89% of interactions less than 1 Mb, whilst in ΔSTAG2 cells, the domains ranged from 160 kb to 6.4 Mb (median 575 kb), with 60% of them less than 1 Mb in the ΔSTAG2 cells **(Figure 2B)**. This represents a shift in the domain size, with decreased numbers of smaller domains (<250 kb), maintenance of intermediate sized domains (250-800 kb) and formation of larger domains (>800 kb) associated with STAG2 loss, which is also visible when comparing the contact maps for STAG2-WT versus ΔSTAG2 HiChIP data (**Figure 2C & S3**).

Using virtual 4C (v4C)[14] plotting enabled changes in chromatin interactions with defined anchor regions between the STAG2-WT and ΔSTAG2 cells to be visualised. We plotted regions where we visually observed altered domain structure between the STAG2-WT and ΔSTAG2 cells in interaction maps and observed large differences in interaction frequency, with the ΔSTAG2 cells displaying greatly reduced interaction frequencies on normalised plots (**Figure 2D and S3**). The loss of STAG2 not only results in a reduced number of domains present, but the domains that are able to form with only STAG1 present predominantly result in the loss of smaller domains and larger domain formation.

### DNA LOOP SIGNAL INTENSITY IS REDUCED WITH CHRONIC STAG2 LOSS

The altered TAD formation observed in ΔSTAG2 cells suggested that loop formation may also be affected in these cells. Therefore, the presence/formation of loops was determined using the Hi-C Computational Unbiased Peak Search (HiCCUPS) tool[15]. The appearance of focal peaks in proximity ligation data is indicative of a “loop” between two DNA locations. The analysis was run at 5 kb, 10 kb and 25 kb resolutions, with the final output being a merged consensus loop list from all 3 resolutions for each replicate library. The final loop call list contained 2,859 loops in the STAG2-WT cells compared to 994 in the ΔSTAG2 cells. Surprisingly, the vast majority of the loops were of similar size range **(Figure 2E)**, albeit it with some loss of smaller loops (<100 kb) in the ΔSTAG2 cells. Although significantly fewer loops were called in the ΔSTAG2 cells, this is not caused by the significant loss of loops of any specific size. Importantly, although much weaker loop peak signals were present in the ΔSTAG2 cells, the formation of new loops was not seen. Aggregate Peak Analysis (APA) of the consensus loop lists showed that despite the low loop peak signal within the ΔSTAG2 cells, clear peak foci were visible, albeit with a reduced pixel signal intensity, when compared to STAG2-WT cells **(Figure 2F)**.

### GENOME COMPARTMENTALISATION ANALYSIS HIGHLIGHTS COMPARTMENT SWITCHING ASSOCIATED WITH CHRONIC LOSS OF STAG2

Transcriptionally active and repressed compartments within the genome are observable within the HiChIP data using Pearson’s correlation matrices of observed over expected read density creating a plaid pattern at a resolution of 50 kb [16]. These patterns can be loosely interpreted as compartments defined as A or B, which correlate with euchromatin and heterochromatin respectively. Eigenvector calculation at the same resolution was used to determine if any differences in compartmental shape and edges occur in the ΔSTAG2 cells compared to their WT counterparts **(Figure 3A)**. Positive eigenvector values were assigned by performing H3K27ac and H3K27me3 ChIP-Seq in both the WT and ΔSTAG2 cells and aligning with the eigenvector output.

**Figure 3:**
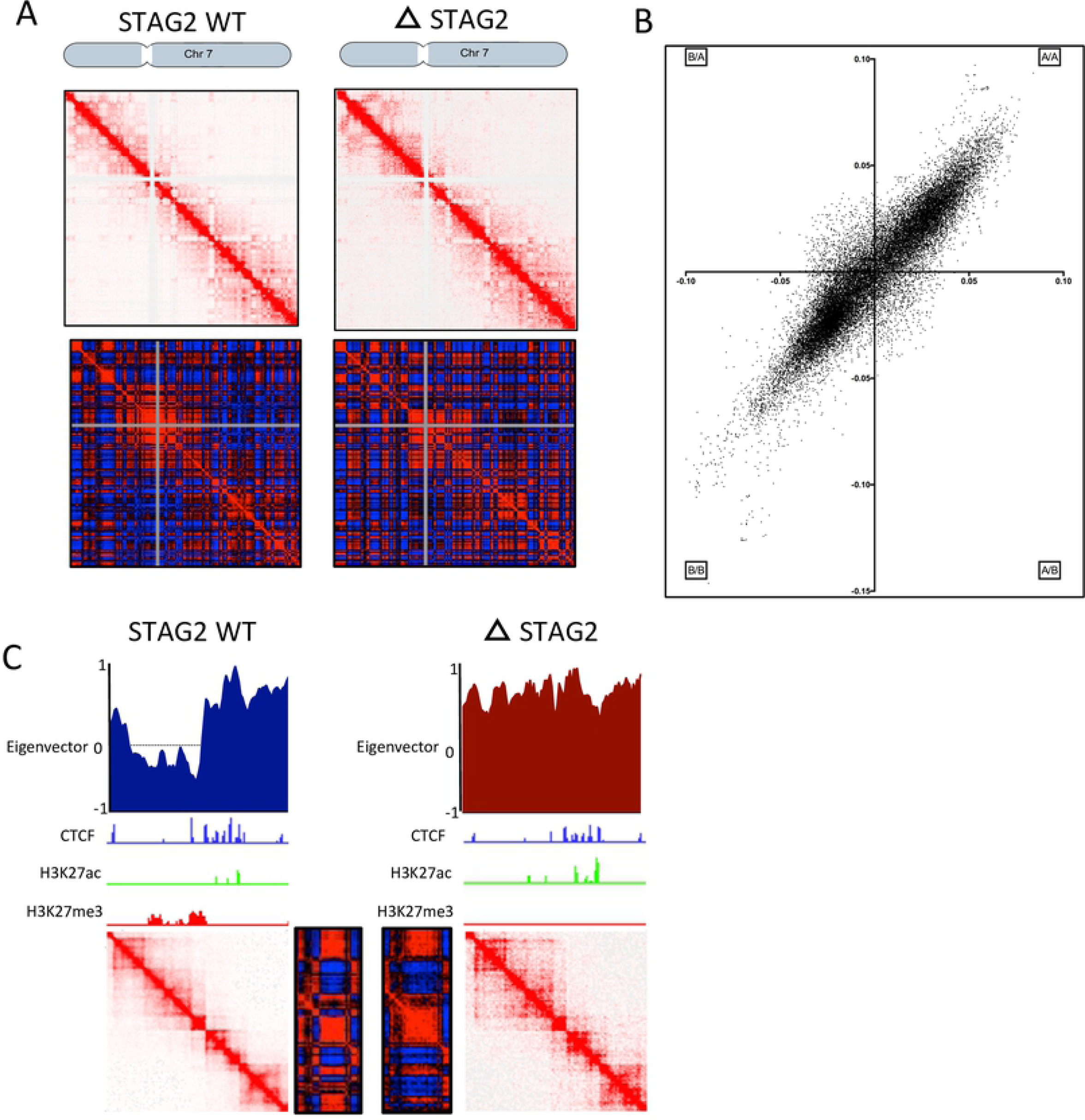
Genome compartmentalization highlights compartment switching. A) Representative HiChIP Contact and Pearson’s matrices of chromosome 7 demonstrating the altered compartmentalisation patterns observed in ΔSTAG2 cells in comparison to STAG2-WT cells. B) Eigenvector compartment scores at 50kb bin sizes highlighting the differences in A/B compartments between WT and ΔSTAG2 cells. C) Example region covering Chr7:20-25mb in WT and ΔSTAG2 cells including; eigenvector scores, HiChIP and Pearson’s correlation matrices and CTCF, H3K27ac and H3K27me3 ChIP-Seq tracks, highlighting the differences in compartment status in STAG2-WT and ΔSTAG2 cells.

Eigenvector scoring of the ΔSTAG2 HiChIP data showed that ∼10% of compartments scored as type B in the WT cells had switched to type A compartments, whilst ∼8% of compartments scored as A were type B compartments in the ΔSTAG2. This “switch” in compartment status is greater than would have been expected by chance (∼5%) (**Figure 3B**).

Furthermore, when comparing the compartmental plaid patterns, ΔSTAG2 compartments appeared to “blend” at edges by switching from A-B or B-A at earlier or later points than seen in the WT counterparts, suggesting a degree of “slippage” at compartment edges (**Figure 3C**).

### CHRONIC LOSS OF STAG2 ALTERS TRANSCRIPTIONAL PROGRAMMES

Previous studies assessing the impact of acute or short-term depletion of members of the cohesin complex showed a modest impact on transcription [17]. To examine the impact of chronic loss of STAG2, a clinically relevant scenario in a relevant genetic background, RNA-seq was performed on the matched isogenic cell pair and the gene expression profiles generated compared **(Figure 4A)**. The chronic loss of STAG2 had a relatively large effect on overall gene expression with 24.5% of the active transcriptome (3149 genes) exhibiting some degree of significant change in expression; 1667 (53%) genes decreased and 1482 (47%) genes increased in expression (**Figure 4B & Table S1**). To identify genes whose expression was more definitively deregulated, a >2 fold-change cut-off identified 225 (32.6%) upregulated and 466 (67.4%) genes downregulated under chronic loss of STAG2 **(Table S1)**. Increasing the cut-off level to >5 fold resulted in the identification of 13 (12.9%) up-regulated and 88 (87.1%) down-regulated genes **(Table S1)**.

**Figure 4:**
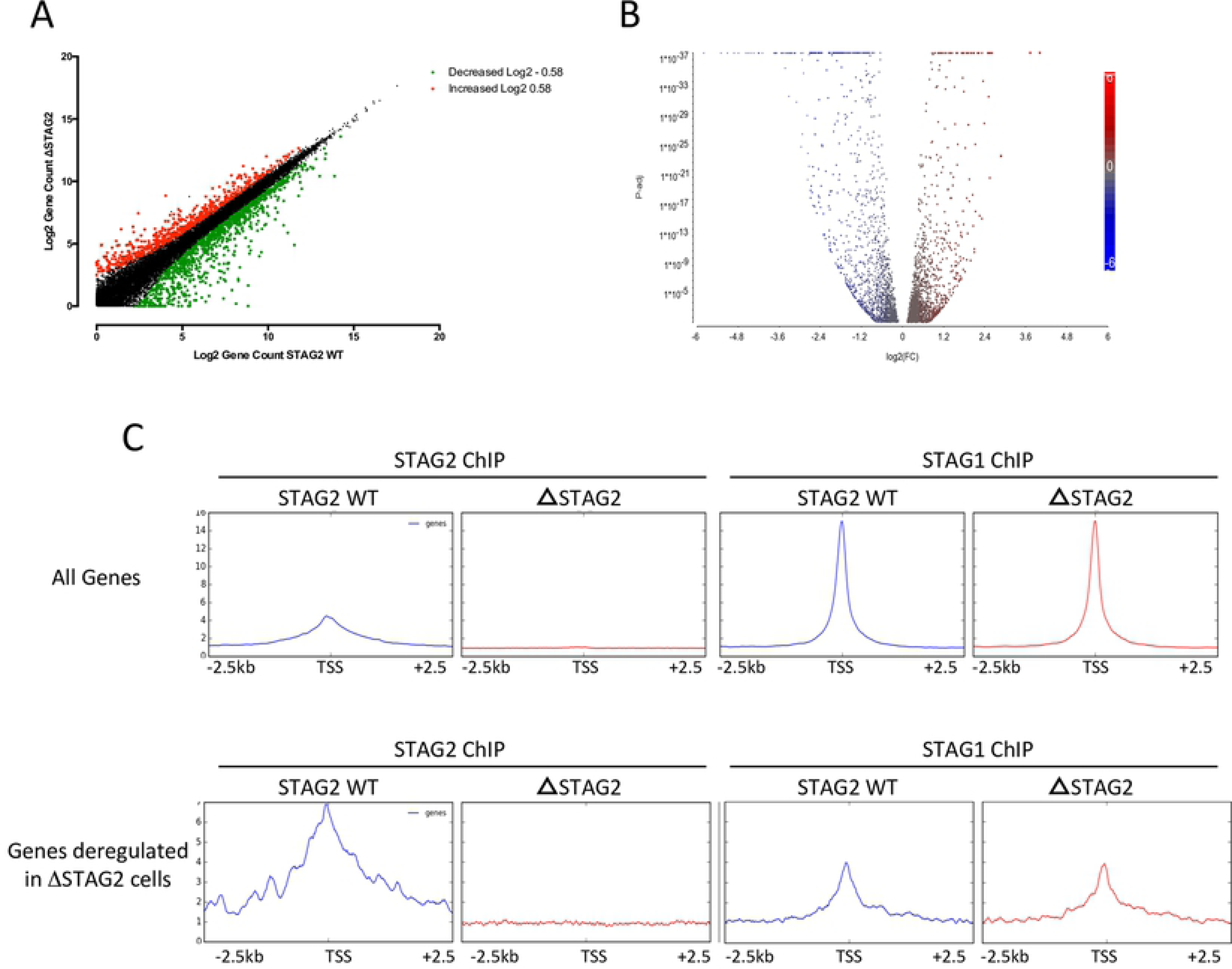
Chronic loss of STAG2 subtly alters global transcriptional programmes. A) RNA-SEQ density plots of Log2 gene counts in the Δ STAG2 cells compared to the WT cells. Changes greater than or less than 1.5 fold (-/+ Log2 0.585) are indicated in green (decreased) and red (increased). B) Volcano plot displaying the 3149 significantly altered (adj p<0.05) genes between the WT and ΔSTAG2 cells. C) Upper Panel - ChIP-Seq generated binding profiles for STAG1 and STAG2 at transcription start sites (TSSs) of all coding genes (Upper Panel) and at the TSSs of genes whose expression was deregulated more than 1.5 fold in the ΔSTAG2 cells (Lower Panel).

To identify specific biological processes affected by chronic STAG2 loss, Ingenuity Pathway Analysis (IPA) of the 691 differentially expressed genes (>2 fold-change) identified key networks that were relevant to “hematopoietic development and maturation”, “cell death” and “cell morphology” (**Figure S4)**. This suggests that whilst the majority of genes and pathways have not become highly deregulated, the chronic loss of STAG2 function has resulted in the transcriptional deregulation of key leukaemogenic pathways, which may contribute to disease development/phenotype.

The disruption in transcription in ΔSTAG2 was unexpected, as previously published data had suggested that STAG1 was predominantly responsible for cohesin binding near gene promoter regions and thereby regulating transcription, whilst STAG2 may be more involved in the maintenance of global 3D chromatin structure [18]. To assess this, the ChIP-Seq data was used to re-analyse STAG1 and STAG2 binding surrounding transcription start sites (TSSs). Average signal density plots confirmed that, on a genome wide level in the STAG2-WT cells, STAG1 was predominantly bound at TSSs in comparison to STAG2 **(Figure 4C)**. Intriguingly, STAG1 binding at TSSs was nearly identical in the ΔSTAG2 cells when compared to the STAG2-WT, suggesting that STAG1 based cohesin complexes contribute to the vast majority of transcription associated interactions in the genome.

Intriguingly, in wild type cells, STAG2 was predominantly bound near the TSSs or promoter regions of the 691 differentially expressed genes than STAG1 (**Figure 4C**). Moreover, STAG1 binding at the TSS did not change in the setting of chronic STAG2 loss, indicating that at these sites, STAG1 was unable to compensate for STAG2. The presence of STAG2, and the lack of STAG1 compensation, at/near the TSSs of these genes further suggest that STAG2 may play a role in linking transcription regulation regions with promoters and enhancers and/or protecting certain promoters from interactions with enhancers in 3-dimensional space.

### GENE EXPRESSION CHANGES MEDIATED BY LOSS OF STAG2 ARE LINKED WITH ALTERED DOMAIN STRUCTURE

To further examine the relationship between STAG2 mediated changes in genome structure and altered transcription, we specifically focused on the *HOXA* cluster, as multiple *HOXA* genes were deregulated upon chronic STAG2 loss, and deregulation of these genes is highly associated with leukaemogenesis [19]. Interestingly, the region containing the HOXA cluster features extensive structural control, with many CTCF/cohesin sites positioned throughout the locus.

From the ChIP-Seq data, seven strong CTCF sites were identified that were associated with the *HOXA* locus **(Figure 5A)**. These sites linked the 3’ region of a TAD (300 kb), to an upstream anchor within the *SKAP2* gene. The major boundary edge of this TAD (termed C7/9 boundary (C=CTCF)) was located between the *HOXA7* and *HOXA9* sites with *HOXA9, HOXA10, HOXA11, HOXA13* and *HOTTIP* lying downstream in a region of high H3K27ac **(Figure 5A)**. The expression of these genes remained relatively unchanged in the ΔSTAG2 cells compared to WT **(Figure 5B)**. In contrast, the genes lying upstream of the C7/9 boundary (*HOXA2, HOXA3, HOXA4, HOXA5, HOXA6* and *HOXA7)* were all significantly up-regulated in the ΔSTAG2 cells (2.2 to 3.41-fold higher than the STAG2-WT cells) (**Figure 5B**). The upregulation of the genes upstream of the C7/9 boundary was also associated with increased H3K27ac and decreased H3K27me3, suggesting an association with altered compartmentalisation. However, the resolution of the Pearson’s correlation and eigenvector calculations used to visualise genomic compartmentalisation is not sufficient to allow us to examine this finite region of the genome. Nonetheless, given that changes in compartmentalisation are generally driven by altered local chromatin structure, v4C was used to visualise the domain region spanning the 300 kb TAD region using the major CTCF boundary site within *SKAP2* as the anchor point **(Figure 5C)**.

**Figure 5:**
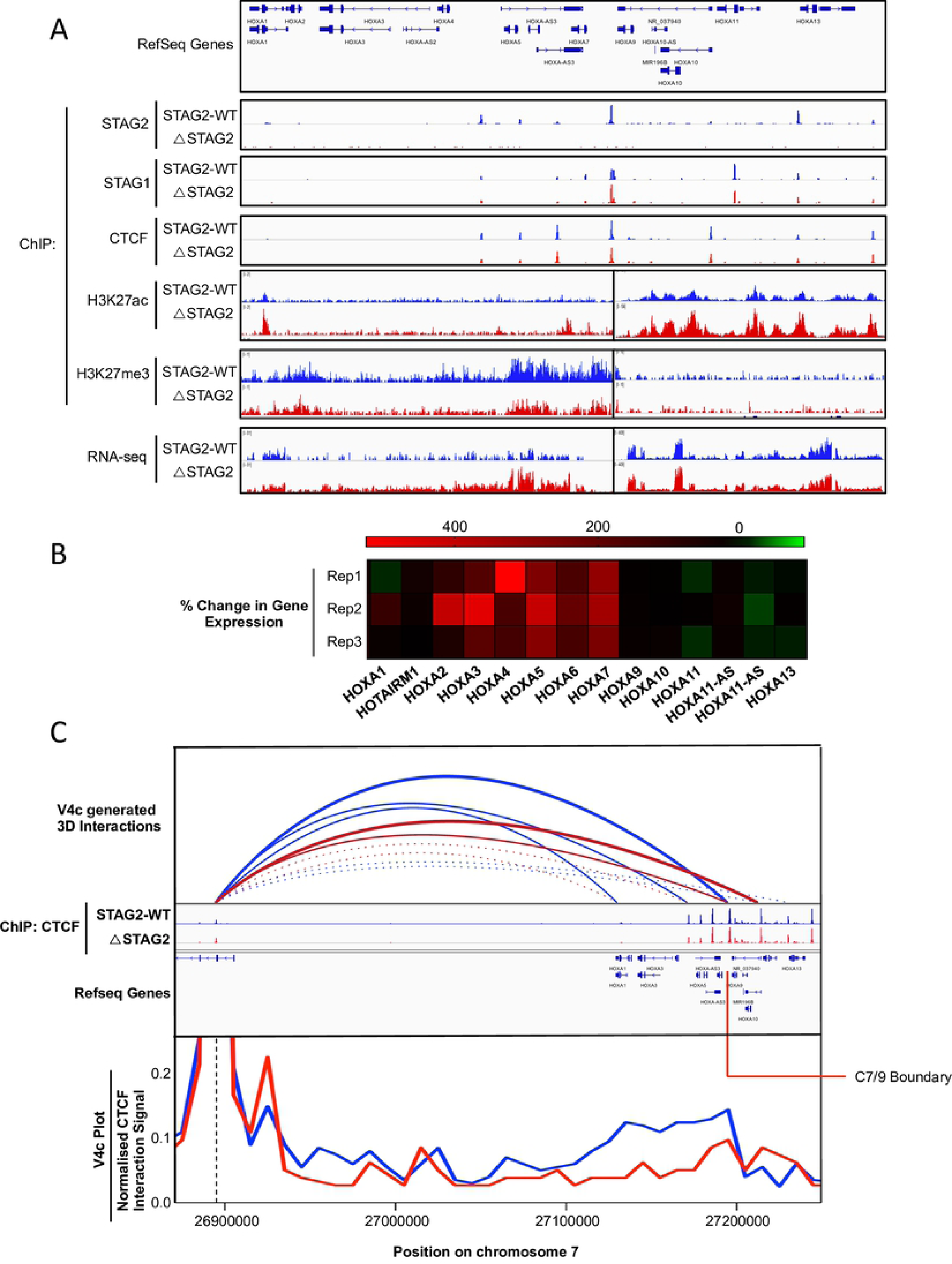
Gene expression changes mediated by loss of STAG2 are linked with altered domain structure. A) ChIP-Seq generated binding profile tracks for STAG2, STAG1 and CTCF across the HOXA cluster region displaying consistent STAG1 and CTCF binding profiles in the absence of STAG2. ChIP-Seq generated H3K27ac and H3K273me tracks show increased H3K27ac across the HoxA locus with decreased H3K273me across the region encompassing *HOXA1-7* in ΔSTAG2 cells compared to STAG2-WT cells. In keeping with this, the normalised RNA-Sew read count track demonstrates increased transcription/gene expression across the *HOXA1-7* region in the Δ STAG2 cells compared to STAG2-WT cells (different scales were used to display H3K27ac, H3K273me and RNA-Seq read count tracks across the *HOXA1-7* and *HOXA9-13* regions in order to highlight the differences in these regions between STAG2-WT and ΔSTAG2 cells. B) Heatmap showing the % change in gene expression between STAG2-WT and ΔSTAG2 cells of all genes located within the HOXA cluster represented above, in each replicate (Rep) RNA-Seq experiment carried out. C) Virtual 4C plot displaying interactions anchored at the CTCF site located within the SKAP2 gene located upstream of the HOXA cluster. CTCF binding tracks along with RefSeq gene tracks have been included for comparison. Additionally, v4c generated interaction loops across the region have been drawn, indicating the reduced interaction frequency between the SKAP2 and the C7/9 CTCF sites, and the increased interaction frequency between the SKAP2 and the C10/11 CTCF sites in ΔSTAG2 cells compared to the STAG2-WT cells. This demonstrates altered 3D structure within the HOXA cluster in ΔSTAG2 cells and indicates the inclusion of early HOX genes (HOXA1-A7 within the transcriptionally active TAD controlling expression of the late HOX genes (HOXA9-13) in these cells.

Significantly reduced interaction frequency and novel points of interaction between the *SKAP2* anchor point and with different CTCF sites within the *HOXA* cluster were identified. The major interaction in STAG2-WT cells occurred between the *SKAP2* anchor point and the C7/9 boundary; the frequency of this interaction was significantly reduced in the ΔSTAG2 cells, with a high-frequency interaction now occurring downstream at the C10/11 boundary, resulting in a larger sub-TAD domain **(Figure 5C)**.

The change in local structure within this TAD may contribute to altered compartmentalisation in this region, as indicated by the altered H3K27ac and H3K27me3 signals across the C7/9 boundary. This would lead to transcriptional activation of the genes within the TAD. Specifically the genes upstream of the C7/9 boundary which would include the early HOX genes (*HOXA2-A7*) would now lie within the transcriptionally active TAD in which the more highly expressed late HOX genes (*HOXA9-13*) are normally placed.

To assess whether the up-regulation of early HOX genes occurs in patients with STAG2 mutant disease, HOXA cluster gene expression was examined in two different public datasets held within the Gene Expression Omnibus (TGCA AML dataset GSE68833 and CD34+ cells from MDS patients, GSE58831) **(Figure S6-S7)** [20]. Although STAG2 mutant disease is rare in these datasets and the fraction of STAG2 mutant cells in either dataset is somewhat diluted due to the presence of non-leukemic lymphoid cells in each sample (the average blast cell count in AML samples is <62%, whilst the average pre-isolated blast cell count is <19%), both datasets showed significant up-regulation of many of the early HOXA genes; A5 and A7 in AML patients and A3, A4 and A7 in MDS patients.

The formation of a larger domain spanning the HOXA cluster led to upregulation of genes normally held within a repressive domain. However, the transcriptional consequences of the loss of smaller domain structures associated with chronic loss of STAG2 were not clear. Interestingly, when examining biologically important pathways from our IPA analysis, a large number of downregulated genes were associated with the MAPK signalling pathway (**Figure 6A-B**). These genes mapped to genomic locations that were associated with loss of small domain structures, which may function to link promoter and enhancer regions (**Figure 6C-D** & **S5A-B**). One gene of interest was DUSP4, which has been linked with sensitivity to MEK inhibition [21]. Analysis of DUSP4 expression in the same publically available datasets described earlier showed that DUSP4 expression was lower, but not significantly, in STAG2 mutant samples compared to STAG2 wild-type samples **(Figure S8)**. This was likely due to the limited number of STAG2 mutated samples, the reduced blast cell fraction in these samples and the more subtle changes in expression of these genes observed in our ΔSTAG2 model.

**Figure 6:**
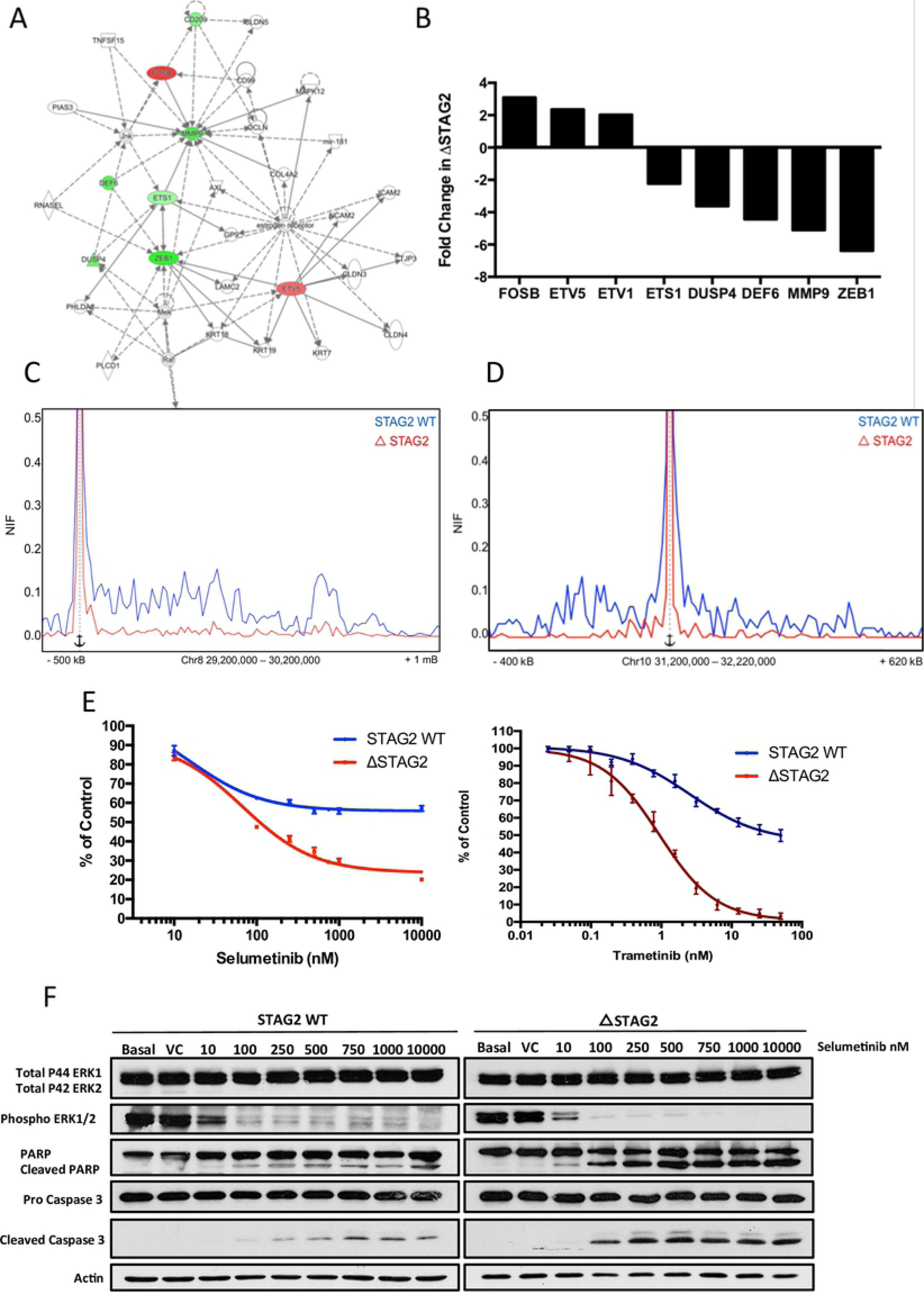
Pathway analysis identifies MAPK signalling as a potential therapeutic target in ΔSTAG2 cells. A) Ingenuity Pathway Analysis (IPA) network diagram displaying altered expression of MAPK signaling related genes in ΔSTAG2 cells. Genes whose expression was increased or decreased in Δ STAG2 cells are highlighted in red and green respectively. B) Fold change in expression of genes specifically identified in the IPA MAPK signaling network generated from normalized RNA-Seq read counts in STGA2-WT and ΔSTAG2 cells C) Virtual 4C plot displaying interactions over a 600 kb region of chromosome 8 encompassing the DUSP4 gene. The V4c plot is anchored upstream of the DUSP4 gene. D) Virtual 4C plot displaying interactions over a 600 kb region of chromosome 8 encompassing the ZEB1 gene. The V4c plot is anchored upstream of the ZEB1 gene. E) Dose response assay displaying increased sensitivity of ΔSTAG2 cells compared to STAG2-WT cells to the MEKi Selumetinib (Left) and Trametinib (Right). Cell death was assessed 24-hours following treatment using the cell titre-glow assay as a readout. Cellular survival in plotted as a percentage signal normalised to control/vehicle treated cells. F) Representative Western Blot showing lower dose Selumetinib mediated inhibition of ERK1/2 phosphorylation and induction of apoptosis (indicated by PARP and Caspase 3 cleavage) in ΔSTAG2 cells compared to the STAG2-WT cells. Densitometry analysis of this blot is shown in Figure S4D-F.

To determine if the downregulation of MAPK signalling genes associated with the loss of smaller domain architecture in ΔSTAG2 cells, the sensitivity of the isogenic cell lines to MEK inhibition was assessed. The ΔSTAG2 cells were significantly more sensitive to treatment with the dual MEK1/2 inhibitors Selumetanib and Trametinib, than the STAG2-WT cells (**Figure 6E**). Consistent with this, ΔSTAG2 cells exhibited lower dose inhibition of MEK1/2 (assessed by ERK1/2 phosphorylation) and extremely low dose induction of apoptosis (assessed by PARP and Caspase 3 cleavage) in comparison to STAG2-WT cells (**Figure 6F & S5C-E**).

## DISCUSSION

How the cohesin complex regulates 3D genome structure has been a focus of intensive study in recent years. However, much of this focus has been on examining the functional role of cohesin in the regulation of 3D chromatin structure through the depletion of loading and release factors, such as NIPBL and WAPL and via transient depletion of the core cohesin component RAD21. Here, we have focused on studying the impact of a clinically relevant mutation in STAG2, a key component of the cohesin complex frequently mutated in MDS and AML. To date, this is the first study examining the function of cohesin following the chronic loss of a single member of the STAG proteins (STAG2) in a clinically relevant genetic background, therefore assessing the potential impact of this mutation on disease pathogenesis. Nonetheless, a number of recent studies have shed light on the functional role of cohesin in the regulation of 3D genome structure. Indeed, Rao et al.[22] recently examined the acute loss of cohesin complex function on chromatin structure using high resolution Hi-C following the degron mediated destruction and restoration of RAD21. This revealed that acute loss of cohesin leads to complete loss of all loop domains across the genome, leading to increased compartmentalisation (due to loss of restrictive loop boundaries), but limited changes to transcription [22]. Similarly, conditional deletion of the Cohesin loading factor NIPBL in mouse liver cells, leads to loss of all loop domain structures across the genome, supporting the important role of cohesin in the formation and maintenance of 3D chromatin structure [23]. In contrast, the deletion of the cohesin release factor WAPL, leads to loss of smaller loop domains and the maintenance and/or formation of larger loop domains [24]. This observation is consistent with the loop extrusion model of loop domain formation, in which cohesin slides along DNA, with DNA loops extruded through the cohesin ring structure until meeting DNA bound CTCF (bound at specific CTCF binding sites), where extrusion is paused/arrested forming a stable loop domain [24]. This loop domain remains stable, until such time as cohesin is released; leading to loss of the loop, or CTCF is released, leading to further extrusion and the formation of a larger loop domain [24]. In the absence of WAPL mediated cohesin release, cohesin residence time on DNA exceeds that of CTCF, resulting in further extrusion and increased loop domain size [24]. Adding to this, a study using single molecule imaging and tracking demonstrated that CTCF has a much shorter chromatin residence time (∼1-2 minute) than cohesin (∼22 minutes), suggesting that cohesin may indeed bypass CTCF sites upon CTCF release/exchange and continue extrusion to form larger loop domain structures [25].

The ChIP-Seq data in our study has demonstrated that in the sustained, chronic, absence of STAG2, STAG1 is able to partially compensate for loss of STAG2 mediated chromatin binding, by redistributing to sites previously and exclusively bound by STAG2. However, the compensatory binding by STAG1 at vacant STAG2 sites was unable to maintain the “normal” 3D structure of STAG2-WT cells and instead lead to the loss of smaller domain structures and the maintenance and formation of larger domain structures, a phenotype strikingly similar to that observed upon deletion of WAPL. We propose two possible scenarios to explain this. One possibility is that by chronically depleting STAG2, we have depleted the pool of free functional cohesin able to be loaded onto chromatin and form loops, the continuous formation and extrusion of which forms smaller domains and sub-domains. Indeed, we did not observe an appreciable increase in STAG1 protein expression in our ΔSTAG2 cells, suggesting that this may indeed be the case. Additionally, the fact that we observed significantly weaker loop signals in STAG2 depleted cells suggests that although not defective, loop formation is significantly reduced, supporting this hypothesis. However, one would predict that this would not lead to the formation of larger domains, but rather simply the loss of small loop domain structures. Nevertheless, it is also possible that STAG1 containing cohesin (STAG1/cohesin), forms more stable DNA bound complexes and/or is more processive (i.e. extrudes DNA through its pore/ring structure faster) than STAG2 containing cohesin (STAG2/cohesin) [17]. In this instance, STAG1/cohesin would remain bound at more distant CTCF sites at the boundaries of large primary TADs, thereby leading to loss of smaller domains and the formation of larger domains in the absence of STAG2/cohesin.

In any event, it is clear that the loss of smaller domains and the generation of larger domains have a wider functional impact in our model system. Indeed, we observed altered genomic compartmentalisation in our ΔSTAG2 cells compared to their STAG2-WT counterparts; compartmental switching occurred at approximately 18% of defined compartments, with slightly more B-A (10%) than A-B (8%) switches occurring. Much of the switching appeared to be the result of “slippage” of compartmental edges, consistent with the formation of larger domains, where adjacent TADs represented the boundary between different compartment types. A similar pattern of slippage at compartmental edges and subsequent compartmental switching was observed in WAPL depleted cells, further linking the formation of larger loop domain structures [24]. The loss of WAPL led to lack of removal of both STAG1/cohesin and STAG2/cohesin and therefore may have an exacerbated effect on genome compartmentalisation. However, this data was generated from HAP1 cells, with a near haploid DNA content, and it is unclear what effect this may have on the gene expression and compartmentalisation requirements of these cells.

A significantly altered gene expression profile was observed in ΔSTAG2 cells as a result of changes in compartmentalisation associated with chronic STAG2 depletion. Although it has been reported that a cancer-associated STAG2 mutation can support sister chromatid cohesion but was unable to repress transcription at DSBs [7]; this may be due to stalled replication fork and collapse of the interaction between the cohesin and the replication machinery [26]

In our study we observed deregulation of the expression of early HOXA genes; aberrant HOXA gene expression is a feature of different sub-types of AML [19, 27]. In the STAG2-WT cells, the HOXA1-7 genes were expressed at very low levels and this region is marked by a low level/absent H3K27ac and a significant level of H3K27me3, which indicates a repressed compartment with the boundary maintained by a TAD lying upstream of the transcriptionally active late HOXA genes (HOAX9-13). However, in the absence of STAG2, the interactions that form and maintain the HOXA1-7 containing TAD are reduced, with formation of a larger TAD containing all of the HOXA1-A10. This new, larger domain also coincided with increased H3K27ac and decreased H3K27me3 across the early HOXA genes (HOXA1-7) region and was coupled with increased expression of these early HOXA genes. During limb development and haematopoiesis, HOX genes are expressed in lineage and stage-specific combinations with gene expression within each HOX cluster generally occurring in a linear fashion, from early to late HOX genes, which allows these genes to “share” enhancers. Most developmental processes are accompanied by two successive waves of HOX gene transcription, with genes in the early and late waves of transcription residing within separate but adjacent TADS [28, 29]. The progression of expression from the early wave of HOX gene expression to late wave of HOX gene expression is controlled by shifting the TAD boundary to allow the “switching” of genes near the TAD boundaries from the TAD containing the early expressed genes, to the TAD containing the late expressed genes [28, 29]. In cells chronically depleted of STAG2 this shift is, at least in part, reversed resulting in increased expression of the early HOXA genes, allowing these genes to interact with enhancers controlling the expression of the late HOXA genes, leading to upregulated expression of this region. The expression of the early HOXA genes may contribute to a block in differentiation, contributing to the development of STAG2 mutated AML. Indeed, the acute siRNA mediated depletion of the core cohesin complex protein RAD21 led to enhanced self-renewal of hematopoietic stem and progenitor cells (HSPCs) driven by de-repression of HOXA7 and HOXA9 in these cells *in vitro* [30].

The altered gene expression across the HOXA cluster was only one example of deregulated gene expression arising from the loss of STAG2. A number of downregulated genes within the MAPK signaling pathway were associated with loss of smaller loop domains in ΔSTAG2 cells. Intriguingly, our ΔSTAG2 cells were highly sensitive to MEK1/2 inhibition, suggesting that the chronic loss of STAG2 and the associated deregulation of MAPK signaling genes had a functional impact on these cells that may be therapeutically targeted [25, 31].

Taken together, our data demonstrates that the chronic loss of STAG2, in a clinically relevant genetic AML background, led to altered 3D genomic structure, causing altered transcriptional programming, which may contribute to disease development, through the upregulation of early HOXA genes. This may present a novel therapeutic targeting strategy, through targeting the deregulated MAPK signalling pathway with MEK inhibitors.

## MATERIALS AND METHODS

### OCI-AML3 and OCI-AML3^ΔSTAG2^ cells

The male OCI-AML3 cell line was sourced from DSMZ (ACC-582). Cells were authenticated using STR profiling at the Genomics Core Technology Unit, Belfast City Hospital prior to model generation. Cells were cultured in RPMI-1640 supplemented with 10% FBS and 100 U/mL penicillin and 100 μg/mL streptomycin at 37 °C and 5% CO2 atmosphere.

ΔSTAG2 cells were generated using lentiviral CRISPR with sgRNA targeting Exon 20 of *STAG2*. Lentivirus was generated using the packaging and envelope vectors psPAX2 and pMD2.G and lentiCRISPR V2 plasmid containing the sgRNA using 293T cells. Viral supernatant was collected, filtered and the target cells transduced by spinoculation at 500 x RCF for 30 minutes. Cells were selected in 1.5 μg/mL puromycin for 7 days before single cell dilution cloning and expansion to isolate clones.

### Western Blotting

1-10^6^ cells were collected during log phase of growth; washed in PBS and lysed using RIPA supplemented with protease inhibitors on ice for 30 minutes. Collected lysates were quantified using BCA assay and 30 μg protein was loaded onto SDS-PAGE gels. Gels were transferred to PVDF membranes and blocked in 5% w/v powdered milk before overnight incubation with the antibody of choice. The following day membranes were washed and incubated with secondary antibodies before chemiluminescent detection was performed.

### Sanger Sequencing

1-10^6^ cells were collected during log phase of growth and lysed using the Zymo quick-gDNA kit as per the manufacturer’s instructions. PCR primers targeting a region spanning the sgRNA target site were used to PCR amplify the edited region in the cell types. PCR products were purified and sent to the Genomics Core Technology Unit at Belfast City Hospital.

### Drug Screening

Cells were plated at 5 x10^5^ cells per mL at a volume of 100 µL in either sterile white (CellTitre Glo) or sterile black (CellTox Green) 96 well culture plates. Cells were exposed to either DMSO vehicle control or increasing concentrations of the experimental compound. DMSO final concentrations did not exceed 0.1%. Following incubation, either 100 µL of CellTitre Glo reagent or 100 µL CellTox Green was added and readout performed as per the manufacturer’s instructions.

### RNA Library Preparation

Total RNA was isolated from 10^6^ cells per biological replicate using the RNeasy kit (Qiagen). Quality was verified using the Agilent RNA 6000 Nano kit on a Bioanalyzer 2100 before moving to library preparation as per the manufacturer’s instructions using the KAPA Stranded RNA-Seq Kit with RiboErase. Libraries were size selected to include fragments 150-700bp in length and sequenced on an Illumina NextSeq 500 using 2 x 75 bp reads.

### ChIP Library Preparation

Between 10^6^ and 20^6^ cells were collected during log phase of growth, washed with PBS and fixed first in 1.5 mM EGS for 18 minutes followed by the addition of methanol-free formaldehyde to a final concentration of 1% for 8 minutes. Glycine was used to quench before cellular lysis. Chromatin was sonicated using a Bioruptor UCD-500 to an average length of 200-500 bp before overnight incubation with the antibody of choice bound to protein G beads. Chromatin bound beads were collected, washed in increasing concentration salt buffer and eluted overnight. Proteinase K was added and samples incubated for 2 hours at 55c before RNase incubation at 37c for 1 hour. DNA was isolated using phenol-chloroform-isoamylalcohol purification and quantified using a Qubit 3.0. Libraries were generated using the Diagenode Microplex V2 library preparation kit as per the manufacturer’s instructions, libraries were size selected to include fragments 150-700bp in length and sequenced on an Illumina NextSeq 500 using 1 x 75 bp read.

### HiChIP Library Preparation

HiChIP was performed as per the published protocol [12]. Up to 10^6^ cells were collected during log phase growth and washed in PBS prior to crosslinking with methanol-free formaldehyde at a final concentration of 1% for 10 minutes. Cells were then lysed in preparation for *in-situ* contact generation. Isolated nuclei were permeabilised, restriction digested with MboI and restriction sites filed with dNTP’s using Biotin-14-dATP. Filled in ends were ligated together at room temperature for 4 hours with rotation before nuclei were lysed, sonicated and the target proteins immunoprecipitated overnight using protein G Dynabeads. ChIP DNA was collected, washed and crosslinks reversed overnight using Proteinase K. ChIP DNA was eluted and samples purified using the DNA Clean and Concentrator columns using double elution steps. The DNA was quantified using a Qubit 3.0 before biotin ligation junction capture using streptavidin C-1 beads. Samples were washed and taken forward for Tn5 Tagmentation. Tagmentation was performed as per the manufacturer’s instructions followed by PCR amplification for 10 cycles. Libraries were size selected to 200-700 bp and sequenced to a shallow depth on the Miseq using 2 x 100 bp reads. Libraries of sufficient quality were then deep sequenced on the Nextseq using 2 x 80 bp or 2 x 43 bp.

The HiChIP, ChIP-Seq and RNA-seq data files are accessible at GEO Series record GSE111537.

### RNA-SEQ data processing

FASTQ files were downloaded from BaseSpace sequencing hub and checked for quality using FASTQC. Reads were aligned to hg19 using STAR and Ensemble gene annotation. Gene count data generated by STAR was used as input to DESeq2 to generate differential expression data as well as normalised count tables.

### ChIP-SEQ data processing

FASTQ files were downloaded from BaseSpace sequencing hub and checked for quality using FASTQC. Reads were aligned to hg19 using Bowtie2 using the “very-sensitive” option. Files were converted from SAM to BAM using SAMTOOLS and indexed. BAM files were converted to BigWig files using the DeepTools package bamCoverage. Reads were normalised during conversion using the “Reads per genome content” option and all downstream analysis was performed on the normalised BigWigs. Peaks were called using MACS3. Peak lists generated were used to generate Heatmaps using the DeepTools package plotHeatmap and density plots using the plotProfile also in DeepTools.

### HiChIP data processing

HiChIP data was processed using the Juicer pipeline [13] following sequencing. The pipeline takes read pairs and aligns to the genome using BWA-MEM. The contact matrixes generated as an output were KR normalised and used in post-processing stages of the pipeline.

### Loops and Domains

Loops were detected using HiCCUPS using default settings at resolutions of 5, 10 and 25 kb creating a merged consensus loop list in both cell types. Domains were called using Arrowhead using default settings at resolutions of 5, 10 and 25 KB with a consensus domain list created. For the ΔSTAG2 cells we found it necessary to scan the contact matrixes manually, overlaying the domain calls from the OCI-AML3 wild type cells and retaining domains observed in both. However, we only retained the domain location and not the score for these manually added domains.

### Compartmentalisation

Compartmentalisation was assessed using the Eigenvector tool in the Juicer pipeline [13]. We generated 50 kb tracks and correlated the A/B compartments with H3K27ac and H3K27me3 ChIP-SEQ data for each cell type using the H3K27ac mark as an indicator of A compartments and the H3K27me3 mark as an indicator of B compartments.

### Changes in transcription

Differences in transcription were assessed by filtering the normalised gene counts generated by DEseq2. Gene counts were converted to Log2 values and any with a count of 0 in both cell lines were removed. A consensus list of genes with expression levels of greater than Log2 count 4 in one cell type was created. This allowed the determination of the core set of transcriptionally active genes in the cells, as well as detecting genes which had dropped below the cut off implemented to define transcription.

## AUTHOR CONTRIBUTIONS

K.I.M and A.T. conceived the project and J.S.S and K.I.M designed all experiments. J.S.S performed model generation, ChIP-Seq, RNA-Seq and HiChIP. J.S.S, K.M.L, C.M.C, K.I.S and K.I.M analysed and interpreted data. S.G.C, F.G.L, S.S.M, K.I.S and K.I.M provided intellectual input. K.I.M & A.T acquired funding. J.S.S, K.I.S and K.I.M wrote the manuscript with input from all authors.

## ACKNOWLEDGEMENTS

This work was supported by a Leukaemia and Lymphoma NI (LLNI) Golden Anniversary Victoria Montgomery Studentship (R2053CNR) and LLNI development grant (R2536CNR). We are grateful to the Genomics Core Technology Unit, CCRCB, in particular Mr Marc-Aurel Fuchs, for access to, and advice using, the Illumina Miseq and Nextseq 500 sequencers and the High Performance Computing Cluster. We would also like to thank all the members of the Blood Cancer Research Laboratory, CCRCB, for insightful comments on experimental progress and the manuscript direction.

---

**Figure S1: Generation of an isogenic model of chronic STAG2 loss.** A) Sanger sequencing chromatograms from OCI-AML3 cells in which the single copy of *STAG2* (X chromosome) was targeted for editing with CRISPR/Cas9 (ΔSTAG2 cells) creating a deletion in exon 20, generating a premature stop codon. Chromatogram from unedited STAG2-WT cells is shown for comparison. B) Normalized RNA-seq gene count levels highlighting the significant reduction in *STAG2* mRNA expression (gene counts include all known STAG2 isoforms) in ΔSTAG2 cells compared to STAG2-WT cells. C) Normalized RNA-seq gene count levels highlighting the mRNA expression levels of members of the cohesin complex in STAG2-WT cells and ΔSTAG2 cells. D) Representative Western Blots demonstrating expression of cohesin complex members in STAG2-WT cells and ΔSTAG2 cells.

**Figure S2: STAG1 compensates for loss of stag2 chromatin binding** Venn diagram of STAG1 binding peaks in STAG2-WT and ΔSTAG2 cells with 14,375 peaks present in both STAG2-WT and ΔSTAG2 cells, but significantly more STAG1 binding peaks in ΔSTAG2 cells compared to STAG2-WT cells.

**Figure S3: Further examples of altered 3D structure within the ΔSTAG2 genome.** A) HiChIP Contact matrixes displaying interactions over a 1 Mb region 29.2 Mb into chromosome 8 (this region encompasses the DUSP4 gene). B) Virtual 4C plot displaying interactions over a 1 Mb region depicted above. C) HiChIP Contact matrixes displaying interactions over a 2 Mb region 60 Mb into chromosome 18 (this region encompasses the BCL2 gene). D) Virtual 4C plot displaying interactions over a 2 Mb region depicted above.

**Figure S4: Haematological developmental processes are deregulated with chronic loss of STAG2** Deregulated Haematological developmental processes gene network identified through Ingenuity Pathway Analysis (IPA) as altered in ΔSTAG2 cells compared to STAG2-WT cells. Genes whose expression was increased or decreased in Δ STAG2 cells are highlighted in red and green respectively.

**Figure S5: Altered chromatin structure surrounding MAPK signaling related genes leads to sensitivity to MEK inhibition.** A) Virtual 4C plot displaying interactions over a 1.2 Mb region of chromosome encompassing the DUSP4 gene. The V4c plot is anchored upstream of the DUSP4 gene. B) Virtual 4C plot displaying interactions over a 1Mb region of chromosome encompassing the MMP9 gene. The V4c plot is anchored upstream of the TSS for MMP9. C-E) Densitometry based quantification of pERK (C), Cleaved PARP (D) and Cleaved Caspase 3 (E), from the Representative Western Blot shown in figure 6G.

**Figure S6: Altered gene expression in an AML patient cohort** Box and whisker plots of gene expression levels (log2) of STAG2 and genes in the HOXA locus between STAG2 mutant patients (n=6) relative to STAG2 wild-type (n=177) AML patients (GSE68833)

**Figure S7: Altered gene expression in an MDS patient cohort** Box and whisker plots of gene expression levels (log2) of STAG2 and genes in the HOXA locus between STAG2 mutant patients (n=6) relative to STAG2 wild-type (n=83) MDS patients (GSE58831)

**Figure S8: Analysis of DUSP4 expression** A) Box and whisker plots of gene expression levels (log2) of DUSP4 between STAG2 mutant patients (n=6) relative to STAG2 wild-type (n=177) AML patients (GSE68833) B) Box and whisker plots of gene expression levels (log2) of DUSP4 between STAG2 mutant patients (n=6) relative to STAG2 wild-type (n=83) MDS patients (GSE58831)

